# Drivers of individual oak tree selection by acorn dispersing animals inferred from a genotyped seedling cohort

**DOI:** 10.1101/559179

**Authors:** Gabriel Gerzabek, Etienne K. Klein, Arndt Hampe

## Abstract

Seed-dispersing animals can strongly influence plant reproductive success and resulting population structures. Few studies have disentangled different drivers of disperser foraging behavior in natural settings and their actual relevance for plant fitness. Here we adopt a novel approach to investigate the drivers of individual trees’ dispersal success in a mixed Pedunculate oak-Pyrenean oak (*Quercus robur* and *Q. pyrenaica*) forest stand. We genotyped a seedling cohort (*n* = 825) upon emergence and performed Bayesian parentage analyses to infer the acorn dispersal success of each oak tree in the stand. We then modeled this estimate as a function of six tree characteristics. The absolute number of animal-dispersed seedlings was exclusively predicted by crop size and the proportion of dispersed seedlings by the number of fruiting oaks in the neighborhood. Neither the oak species nor tree height, acorn size or shape played any role. Our findings contrast with results from experimental studies and suggest that effective acorn dispersers, despite being scatter-hoarders, behaved much like avian dispersers of fleshy-fruited species when selecting trees to forage on. Their behavior should favor the dominance of large, prolific trees for the dynamics and genetic composition of naturally regenerating oak stands.

## INTRODUCTION

Seed dispersal by vertebrates is considered a key innovation in angiosperm evolution (Tiffney 2004). Effective seed dispersal is critical for the successful establishment of new individuals and the resulting dynamics of plant populations and communities (Schupp et al. 2010). Seed-dispersing animals hence can exert a direct and potentially strong influence on the fitness of reproducing plants through their choice of the individuals they forage on. Their activity leaves measurable imprints in the genetic structure and diversity of plant populations (García and Grivet 2011, Hamrick and Trapnell 2011) and can drive rapid microevolutionary changes (Galetti et al. 2013). A better understanding of the cues that guide foraging seed dispersers in their choice of fruiting plants hence can provide valuable insights into the role of plant-disperser interactions for the reproduction and resulting demo-genetic structure of plant populations. Yet few studies have achieved to disentangle and compare different drivers of seed disperser behavior under natural conditions. Moreover, short-term observations or experiments tell little about the actual relevance of disperser behavior for plant reproductive success.

Seed dispersers rely on a complex system of decision cues and spatial memory for selecting the plants they forage on (Corlett 2011). Decision cues can be used hierarchically or sequentially, and their role can vary depending on the animal’s spatial scale of perception or nutritional status. Scatter-hoarding species that collect and store seeds for later consumption tend to behave differently than species that seek fleshy fruits to eat them immediately (Vander Wall and Beck 2012). Most studies on disperser foraging decisions have focused on cues related with either fruit abundance or fruit traits. Fruit crop size usually is a consistent predictor for the amount of seeds removed from individual plants (Carlo and Morales 2008, Prasad and Sukumar 2010). The abundance of fruits in the neighborhood (conspecific or heterospecific) can also influence rates of seed removal, either positively through an increased attraction of dispersers (Morales et al. 2012) or negatively owing to their satiation (Saracco et al. 2005, Hampe 2008). On the other hand, diverse morphological, physical and chemical fruit traits have experimentally been shown to influence fruit choice, highlighting the capacity of frugivores to differentiate even subtle signals (Corlett 2011). Fruit traits can also influence disperser foraging behavior under natural conditions, although their actual relevance varies greatly across systems (Jordano 2000, Vander Wall 2010).

Despite these advances, our understanding of disperser behavior and its consequences for plant recruitment remains fragmentary. Studies on scatter-hoarders have largely focused on fruit characteristics but rarely on the identity of the source plant (Vander Wall 2010, Pesendorfer et al. 2016; but see Wästljung 1989, Morán-López et al. 2015), while studies of other frugivores have focused much on fruit abundance and less on individual-level variation in fruit traits (but see e.g. Sallabanks 1993, García et al. 2001). Most importantly, virtually no studies have to date linked frugivore behavior with the fitness of the fruiting plant due to the difficulty of tracking animal-mediated seed dispersal events and resulting plant establishment (Schupp et al. 2010). Molecular ecological approaches can help overcome this limitation (García and Grivet 2011). Parentage analyses based on genetic markers allow to infer the source plants of dispersed seeds or seedlings and thus to quantify the reproductive success of all adults in terms of the number of descendants they have achieved to disseminate and establish across the population. This fitness measure can then be related with characteristics of the fruiting plant and its environment.

Here, we combine field and molecular ecological research approaches to assess drivers of individual oak tree selection by acorn-dispersing animals in a mixed Pedunculate oak-Pyrenean oak (*Quercus robur* and *Q. pyrenaica*) forest stand. Effective acorn dispersal is primarily performed by the Eurasian jay (*Garrulus glandarius*) in our study area (Gerzabek et al. 2017). Jays are major dispersers of oak acorns across much of temperate Eurasia (Pesendorfer et al. 2016). They are very efficient scatter-hoarders that can harvest, disperse and store thousands of acorns within a single fruiting season (Kollmann and Schill 1996, den Ouden et al. 2005). Their foraging behavior hence can have direct and significant implications for the relative contribution of individual oak trees to plant recruitment at forest stand level (Pesendorfer et al. 2018). Eurasian jays have been shown to favor large and slim acorns and to differentiate among oak species in choice experiments (Bossema 1979, Pons and Pausas 2007, Myczko et al. 2014). But it remains little known whether their foraging decisions under natural conditions are primarily guided by acorn traits, local acorn abundance, or other drivers (Pesendorfer et al. 2016, Morán-López et al. 2015).

We genotyped a cohort of recently germinated oak seedlings and performed parentage analyses to quantify the effective dispersal success – estimated through the absolute number or the proportion of seedlings emerging away from the mother tree – of each adult oak growing within the forest stand. This estimate was then related with several tree features to simultaneously test how these influenced the choice of individual oak trees by foraging jays. Specifically, we addressed the following research questions: 1) How many seedlings does each tree of the forest stand contribute to the genotyped cohort? 2) Can trees’ dispersal success be predicted from their species identity, height, crop size, acorn size or shape, or oak abundance in their neighborhood? 3) What is the relative importance of these predictors for explaining acorn dispersal success?

## MATERIALS AND METHODS

### Study system

The study was performed in the Forêt de Nezer (44°34’ N, 1°00’ W) ca. 50 km SW of Bordeaux, SW France. The area is covered by extensive plantations of maritime pine (*Pinus pinaster* Ait.) interspersed with small stands of broadleaf forests dominated by Pedunculate oak (*Quercus robur* L.) and, to a lesser extent, Pyrenean oak (*Q.pyrenaica* Willd.). Such stands are largely exempt from forest management. We selected a mixed oak forest stand with ca. 280 adult Pedunculate and Pyrenean oaks (90% *Q. robur*, 10% *Q. pyrenaica*) for this study. A detailed description of the stand can be found in Hampe et al. (2010). Acorn dispersal in the area is performed by Eurasian jays as well as by rodents (primarily wood mice, *Apodemus sylvaticus* L.) (Gerzabek 2016). Importantly, several jays were regularly present in our study stand and different lines of evidence – field observations, the spatial locations and microhabitats of sampled seedlings (Gerzabek 2016, Gerzabek et al. 2017) and complementary experiments on acorn predation and dispersal on the ground (A. Hampe, unpublished data) – suggest that only a limited proportion of the seedlings we sampled would have been dispersed and cached by other animals than jays.

### Field sampling and laboratory analyses

In early spring 2006, we delimited a study plot of 6 ha enclosing the oak forest stand and surveyed all adult trees within this area and an adjacent belt of 100 m width. Trees were identified as adults based on their size and on fruit set observations. Each tree was individually tagged and mapped, its diameter at breast height (dbh) was measured, and several buds were collected and stored at −80° C for later DNA isolation. We subsequently measured six further variables for a subset of 79 arbitrarily chosen trees: (i) The oak species (*Q. robur* or *Q. pyrenaica*) was determined in the field based on leaf morphology. (ii) Tree height was measured with an ultrasound vortex. (iii) Acorn crop size during the 2005 fruiting episode was quantified by counting the number of empty acorn cups in four 50×50 cm squares randomly placed beneath the canopy in early 2006 and multiplying the census mean with the dbh of the target tree (which served as a proxy for the canopy size). Moreover, we collected 50 acorns from each tree in September 2006 to determine (iv) acorn size (by measuring their weight to the nearest 0.1 mg) and (v) acorn shape (by measuring length and width to the nearest 0.1 mm and calculating their ratio). (vi) We quantified the abundance of adult oaks in the neighborhood by summing the number of trees within a radius of 10 meters around the focal tree.

During late April and early May 2006, we performed a comprehensive survey of newly emerged oak seedlings (i.e., those stemming from the 2005 fruiting episode) in our 6 ha study plot. We sampled all seedlings emerging more than 2 m away from any adult oaks, searching every part of the study plot at least twice and up to six times. Furthermore, we sampled 20% of all seedlings emerging beneath adult oaks (i.e., up to 2 m beyond the canopy projection to account for potential short-distance dispersal of acorns by wind or bouncing off of branches). These 20% were sampled using a strict randomization protocol to ensure their representative character (see Hampe et al. 2010 for details). We estimate that the resulting data set (*n* = 825 individuals) includes 25-30 % of the overall seedling cohort within the study plot. All sampled seedlings were individually tagged and mapped, and one leaf was collected and stored at −80°C until DNA isolation.

Trees and seedlings were genotyped using eight nuclear microsatellite (SSR) markers (QrZAG11, QrZAG96, QrZAG112, QpZAG110, QrZAG5b, QrZAG7, QrZAG20, and QrZAG87) as described in detail in Hampe et al. (2010). In addition, we genotyped all individuals at 39 SNP loci as described in Gerzabek et al. (2017). We obtained readily usable data for 33 SNPs that we merged with the SSR data to obtain individual multilocus genotypes. Both the genotypic data and the field data underlying this study have been deposited in Dryad (doi:10.5061/dryad.3j33t).

### Data analyses

Our analysis proceeded in two steps. First, we quantified the fecundity and acorn dispersal success of individual trees. Then we assessed how dispersal success was affected by a series of tree-related parameters.

#### Quantifying individual dispersal success

We performed a parentage analysis using a Bayesian approach as implemented in the software MEMMseedlings (Gerzabek et al. 2017). Unlike many other parentage assignment procedures (but see Moran and Clark 2011), our approach jointly considers genotypes, spatial locations and tree mating parameters to infer parentage and allowed us to distinguish between the mother and the father tree of a given seedling – a highly relevant information for our study purpose. We adapted MEMMseedlings to use a two-step procedure (explained in detail in Gerzabek et al. 2017) where the first step consisted in the estimation of all fecundity and dispersal parameters. The second step consisted in computing the posterior probability of each tree within the stand to be the mother of a given sampled seedling and the posterior probability that this seedling originates from seed migration. Based on these probabilities, we then assigned each sampled seedling categorically to its most likely mother tree (that is, either the tree with the highest posterior probability or a non-sampled mother-tree from outside the stand if the posterior probability of immigration exceeded those of the local mother trees).

Our parentage analysis served two purposes: First, it enabled us to identify the mother trees of our seedlings and to estimate the maternal reproductive success of each tree within the stand by summing up the number of seedlings assigned to this individual. Second, it allowed us to infer categorically whether a given seedling had been dispersed away from its source tree or not (we consider that an animal is always the vector in the former case). We considered that any seedling growing beneath its assigned mother tree stemmed from an acorn that had directly fallen from the mother tree to the ground without being handled by an animal seed disperser. These acorns were classified as “non-dispersed” and the remaining acorns (i.e., those growing either away from any adult oak or beneath an oak that was not assigned as their mother tree) as “dispersed”. The sum of dispersed acorns was used as a proxy for individual trees’ “absolute” dispersal success and the proportion between the dispersed acorns and the overall number of acorns mothered by a given tree was used as an estimate of its “relative” dispersal success.

#### Predictors of individual dispersal success

We constructed generalized linear models to assess the relative effect of different tree features on the absolute and the relative dispersal success of individual trees. Models included the following predictor variables: tree species, tree height, crop size, acorn size, acorn shape, and adult oak abundance in the neighborhood. Crop size and adult oak abundance were log transformed to reach a better fit. We considered no interaction terms in the final model after conducting preliminary tests. Our models assumed quasi-Poisson (absolute dispersal success) and binomial (relative dispersal success) errors.

Finally, we assessed direct relationships between tree crop size and absolute dispersal success using an analysis of density dependence. For this purpose, we performed a linear regression with our crop size index as predictor and the number of successfully dispersed seedlings as response variable (both log-transformed). All but the parentage analysis were performed in R version 3.3.0 (R Development Core Team 2016).

## RESULTS

We could assign a total of 678 seedlings (82% of the overall sample) to a local mother tree and classify 307 of them as non-dispersed and 371 as dispersed. A total of 115 trees (39% of the local adult population) mothered at least one seedling and 87 (30%) at least one successfully dispersed seedling. The distribution of seedling production was highly skewed (Fig. 1), although the trend was weaker among the dispersed seedlings (Kolmogorov-Smirnov: *D* = 0.24, *P* = 0.002).

**Figure 1.**
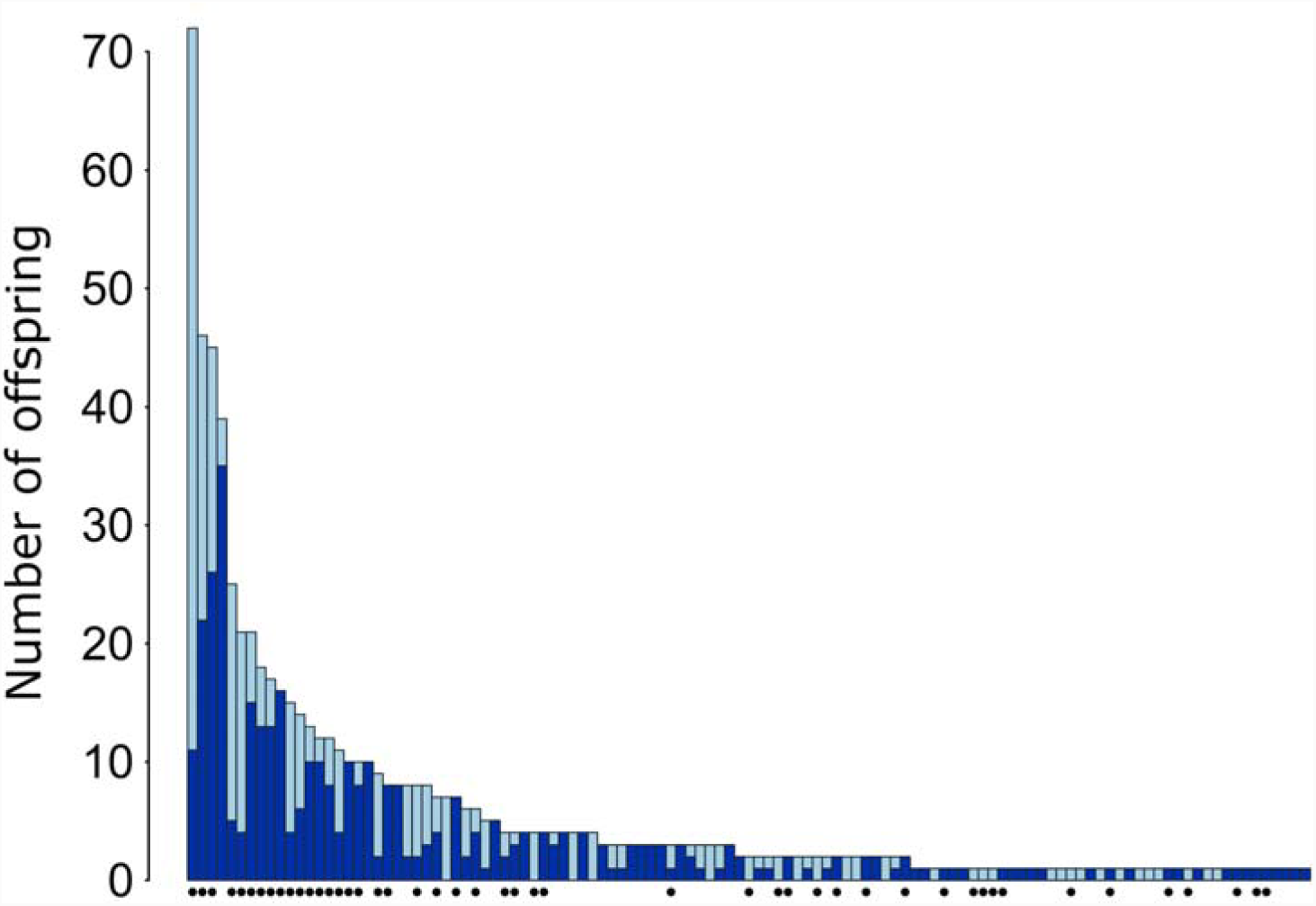
Seedling production by individual *Quercus robur* mother trees. Each column corresponds to a reproducing tree. Dark blue bars indicate those seedlings that were actually dispersed away from the mother tree and light blue bars those that failed to be dispersed. Black circles beneath columns indicate trees for which individual trait data were gathered.

The subset of 79 trees that we had characterized in the field produced 468 seedlings, of which 237 had been dispersed. A total of 47 trees mothered at least one seedling and 38 at least one successfully dispersed seedling.

### Predictors of individual seed dispersal success

Most tree-related variables showed large variation (Table S1 in the Online Appendix). The generalized linear model on absolute dispersal success revealed a marked positive effect of crop size (slope = 0.78), whereas relative dispersal success was exclusively predicted by, and negatively related (slope = −1.26) with, the number of fruiting oak trees within the neighborhood of the focal tree (Table 1; Figures S1a and S1b in the Online Appendix show the underlying relationships between absolute and relative dispersal success with the different predictor variables). Figure 2 shows the observed values and model predictions for the two significant variables (while Figure S2 in the Online Appendix shows predictions for the same model with intervals estimated by 10,000 permutations). Neither the oak species nor tree height or the acorn traits size and shape exerted any noteworthy effect on dispersal success (either absolute or relative). Finally, the regression between our crop size index and absolute dispersal success generated a slope of 0.30 (*F* = 29.7, *P* < 0.001, *R*^2^ = 0.28), significantly smaller than 1 (*t* = −12.52, df = 77, *P* < 0.001).

**Table 1.** Effects of the oak species, tree height, crop size, acorn size and shape, and the number of adult oaks in a radius of 10 meters around the target tree on the absolute number (= absolute dispersal success) and the proportion (= relative dispersal success) of effectively dispersed offspring according to generalized linear models (type II ANOVA, Wald χ^2^ test).

| Variable | Absolute dispersal success |  |  | Relative dispersal success |  |  |
| --- | --- | --- | --- | --- | --- | --- |
| | *F* | d.f. | *P* | $\chi^2$ | d.f. | *P* |
| Species | 1.43 | 1 | 0.23 | 1.24 | 1 | 0.27 |
| Tree height | 0.21 | 1 | 0.64 | 0.38 | 1 | 0.54 |
| Crop Size | 25.87 | 1 | **<<0.001** | 0.23 | 1 | 0.63 |
| Acorn size | 1.27 | 1 | 0.26 | 0.77 | 1 | 0.38 |
| Acorn shape | 0.66 | 1 | 0.42 | 0.95 | 1 | 0.33 |
| Number of adult oaks | 1.13 | 1 | 0.29 | 48.72 | 1 | **<<0.001** |

**Figure 2.**
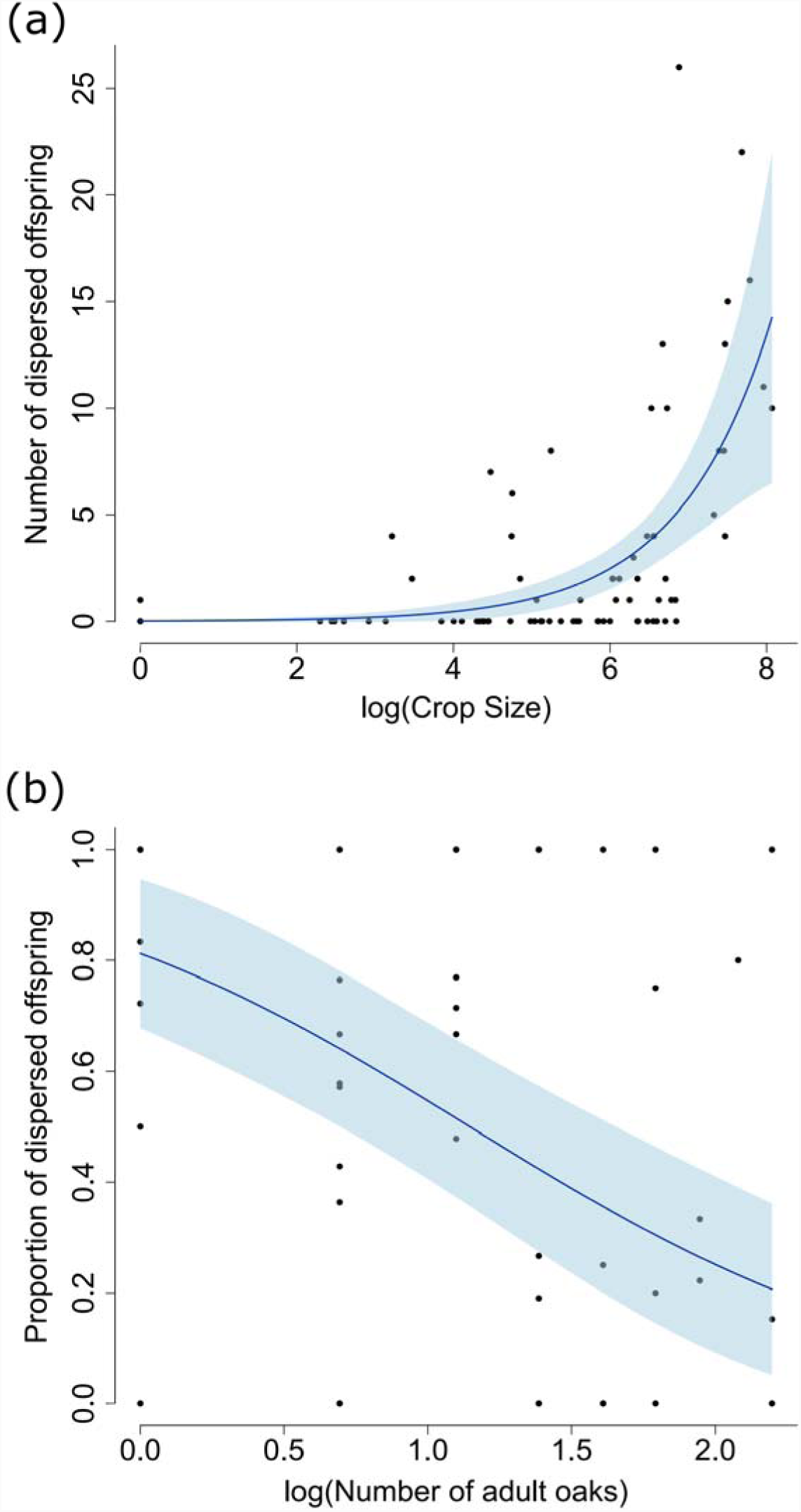
Observed values and model predictions for each of the two single significant predictors that we identified in our regression analyses of absolute (a) and relative (b) acorn dispersal success. Shaded areas indicate the 95% confidence intervals.

## DISCUSSION

### Dispersal success of individual trees

A large part of the seedling cohort stemmed from only a handful of mother trees while many trees did not at all contribute to the seedling pool. Strong inequality in individual fecundity is a widespread phenomenon in tree populations (Moran and Clark 2012) and can persist into the stage of dispersed and established recruits (Schnabel et al. 1998, Sezen et al. 2007, Hampe et al. 2013). Our systematic distinction of dispersed and non-dispersed offspring enabled us however to detect that the skew in tree fecundity was weaker among the dispersed descendants than among the overall cohort. The most likely explanation for this difference is that trees with large crop sizes have a lower proportion of their acorns dispersed as a consequence of disperser satiation (Hampe 2008). The existence of such an effect is confirmed by the analysis of density dependence, and it is little surprising: Even highly efficient scatter-hoarders such as Eurasian jays (Bossema 1979, Kollmann and Schill 1996, Pesendorfer et al. 2016) should be overwhelmed by a stand-scale crop that is likely to sum hundreds of thousands of acorns. Our result is at odds with the hypothesis proposed by Vander Wall & Beck (2012) that scatter-hoarding seed dispersers should rarely be satiated when foraging (but rather when recovering the cached seeds). The rather marked satiation effect that we observed (regression slope between crop size and dispersal success = 0.30) implies that acorn dispersers are a limited resource in our system for which fruiting trees have to compete. Under such circumstances, tree features that influence disperser behavior can directly affect dispersal success and resulting mother tree fitness (Gerzabek et al. 2017).

### Tree-related predictors of dispersal success

Both absolute and relative dispersal success were exclusively driven by patterns of acorn abundance, although in different ways. The most fecund trees achieved to effectively disperse the greatest absolute amount of descendants. This finding fully supports the predator dispersal hypothesis (Vander Wall 2010, Pesendorfer et al. 2016), which poses that larger seed crops result in fitness benefits from increased seed dispersal by seed-hoarding animals. When we accounted for the effect of tree fecundity by calculating relative instead of absolute dispersal success, we found that this measure was negatively related with the abundance of adult oaks in the neighborhood of the target tree. Although we could not quantify the crop sizes of all these trees, the number of adult oaks is likely to represent a reasonably suitable proxy for the local abundance of acorns. In line with our detection of a satiation effect, this finding indicates that trees are competing for dispersers with their direct neighbors (corresponding to the spatial scale addressed by our 10-m radius).

On the contrary, neither the oak species nor acorn size or shape had any measurable influence on dispersal success (although the first result must be interpreted with some caution given the absence of highly fecund *Q. pyrenaica* individuals; see Figs. S1a and b in the Online Appendix). Our finding is noteworthy because to date most quantitative studies of drivers assumed to influence the behavior of acorn dispersers have primarily focused on the oak species as well as morphological or chemical traits of the acorns (e.g. Bossema 1979, Scarlett and Smith 1991, Pons and Pausas 2007, Wang and Chen 2009, Shimada et al. 2015). These traits have often been shown to influence the choice of scatter-hoarders under experimental conditions. However, their actual relevance for effective acorn dispersal success in natural settings is questioned by the results of our multi-factor study, which confirm instead the role of more rarely investigated drivers such as the crop size or the spatial distribution of fruiting trees (Morán-López et al. 2015, Pesendorfer and Koenig 2016; see also Pesendorfer et al. 2016).

Taken together, our results suggest that the selection of trees by acorn dispersers relies primarily on visual choice based on acorn abundance and not on features that can only be properly perceived from within the tree (such as acorn traits) and would have to be memorized until subsequent foraging visits. In this sense, acorn dispersers in our system behaved much like many avian frugivores of fleshy-fruited species (Saracco et al. 2005, Hampe 2008, Carlo and Morales 2008, Prasad and Sukumar 2010, Vander Wall and Beck 2012). Our research approach prevented us from assessing whether acorn dispersers, once settled within or beneath a given tree, tended to select certain acorn phenotypes over others. Such hierarchical selection behavior is also known from frugivores of fleshy-fruited species (Sallabanks 1993, Corlett 2011). However, acorn choice within the same tree should be of relatively minor relevance for its dispersal success and resulting fitness.

### Consequences for oak regeneration

Foraging decisions of seed dispersers are particularly important for plant species that depend on a small suite of vectors for their dispersal. This is the case with the European oaks, for which the Eurasian jay is widely acknowledged to be the major effective seed disperser (Bossema 1979, Gómez 2003, Pesendorfer et al. 2016). The species’ relevance stems not only from the quantity of acorns it can mobilize and the average distance of dispersal events but also from its tendency to increase the survival chances of dispersed acorns by hiding them within the ground, often in microhabitats that are favorable for seedling establishment (Bossema 1979, Kollmann and Schill 1996, Gómez 2003, den Ouden et al. 2005). Different lines of evidence suggest that jays are likely to be responsible for much of the effective acorn dispersal in our study stand (Gerzabek et al. 2017). Our findings that they (i) base their foraging decisions on tree crop size and (ii) are satiated by the stand-scale acorn crop implies that the foraging behavior of jays tends to favor the dominance of large, prolific trees in the recruitment and resulting genetic composition of naturally regenerating oak forests. A previous study performed in our oak stand revealed that the spatial genetic structure of the adult tree population strikingly resembles that of the dispersed seedling cohort (Hampe et al. 2010). This similarity was explained by the fact that most younger adults and most dispersed seedlings probably stem from the same few large founder trees. If true, it would represent an empirical evidence that the foraging behavior of acorn dispersers such as jays may indeed have a measurable long-term impact on the population structure of naturally regenerating oak forests.

## ACKNOWLEDGEMENTS

We thank Begoña Garrido and Jean-Marc Louvet for their excellent support during field work and Sylvie Oddou-Muratorio for her help with MEMMseedlings. The study was funded by a doctoral grant from Bordeaux University to G.G., the INRA ACCAF project FORADAPT and the projects ExpandTree (ANR-13-ISV7-0003) and SPONFOREST (BiodivERsA3-2015-58). A.H. conceived the study, G.G. and A.H. performed the field and laboratory work, and G.G., E.K.K. and A.H. analyzed the data. All three co-authors contributed to the final text and approved its final version.

