## Supplementary material for "Drivers of individual oak tree selection by acorn dispersing animals inferred from a genotyped seedling cohort": Online Appendix

**Table S1**. Individual variation in different traits measured at 79 adult oak trees from the target forest stand. Note that trees were arbitrarily chosen and values hence are not fully representative of the stand.

| Variable | Mean ± SD | Range |
|---|---|---|
| Crop size index | 553 ± 695 | 0 - 3194 |
| Tree height (m) | 15.3 ± 3.7 | 9.2 - 25.0 |
| Acorn weight (mg) | 3962.3 ± 1075.8 | 1822.5 - 7374.3 |
| Acorn shape (ratio length:width) | 1.7 ± 0.2 | 1.2 - 2.3 |
| Number of adult oaks within 10m | 2.8 ± 2.3 | 0 - 8 |

**Figure S1a.** Graphical representation of the individual relationships between absolute dispersal success (measured as the absolute number of dispersed seedlings per mother tree) and the different mother tree related predictor variables. Oak species: P = *Quercus pyrenaica*, R = *Quercus robur*.

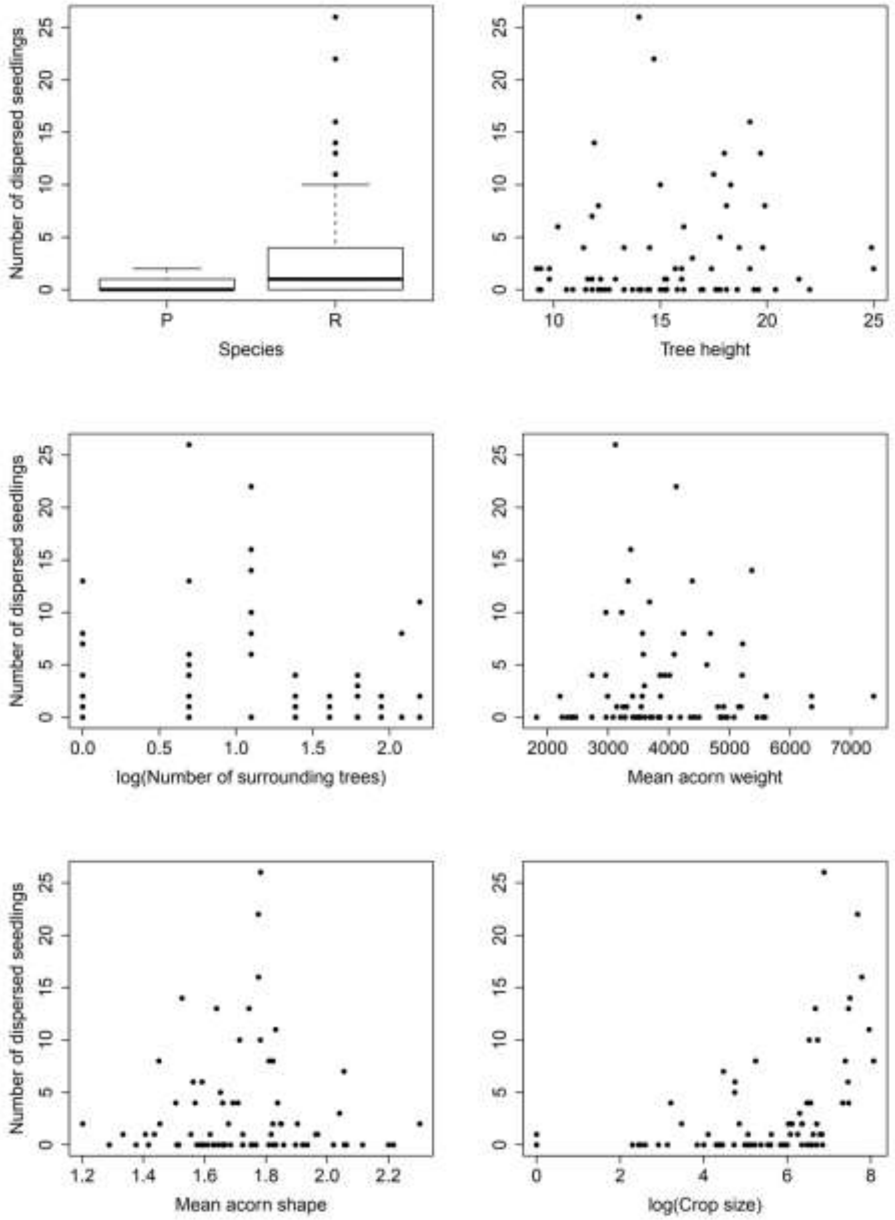

**Figure S1b.** Graphical representation of individual relationships between relative dispersal success (measured as the proportion of seedlings from a given mother tree that had been dispersed) and the different mother tree related predictor variables. Oak species: P = *Quercus pyrenaica*, R = *Quercus robur*.

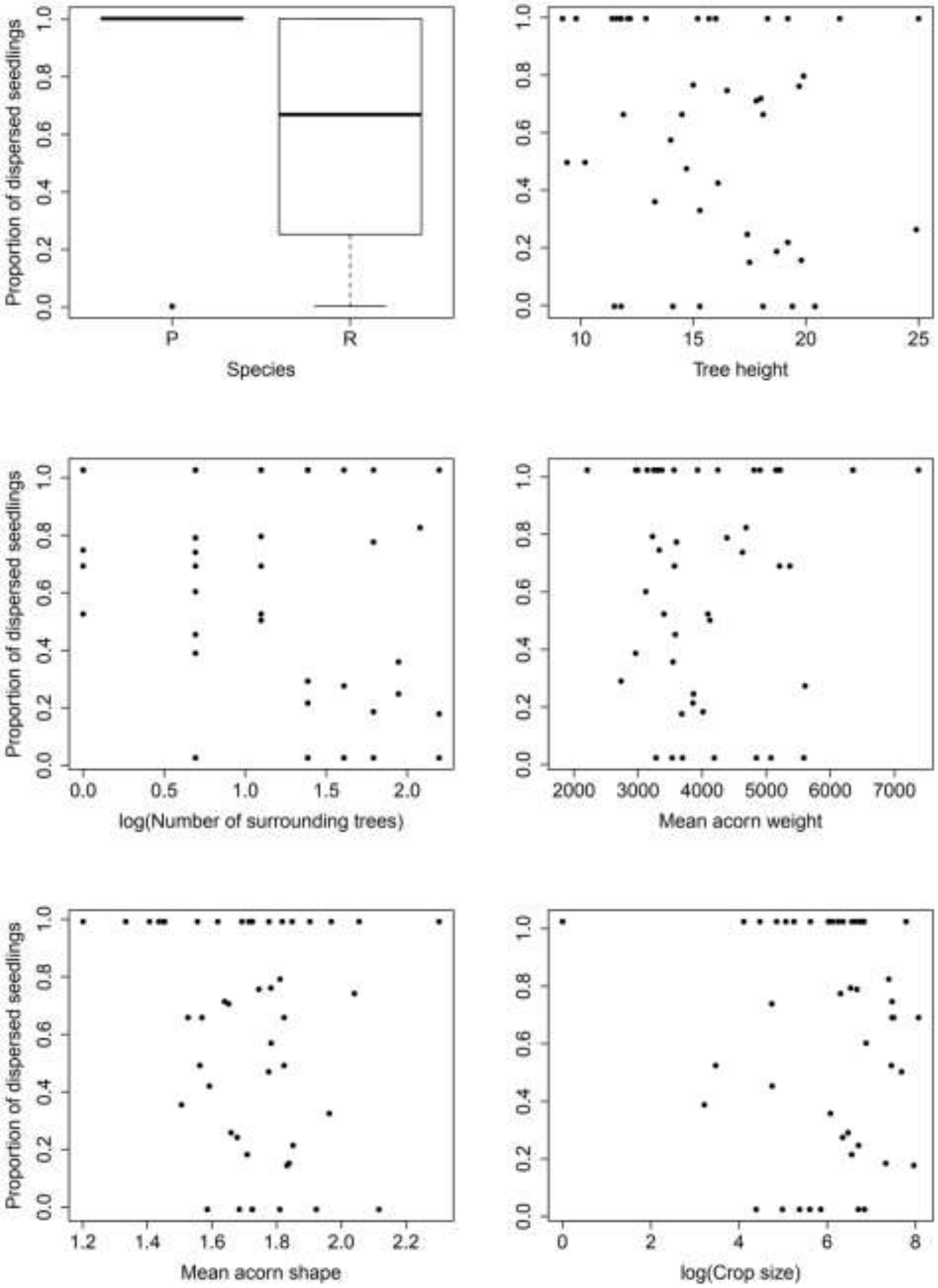

**Figure S2.** Predicted effect of crop size on absolute dispersal success for the full model as shown in Figure 2 but with prediction intervals estimated by a bootstrap with 10,000 permutations. Note that drawing the same plot for the binomially distributed response variable relative dispersal success would not be informative.

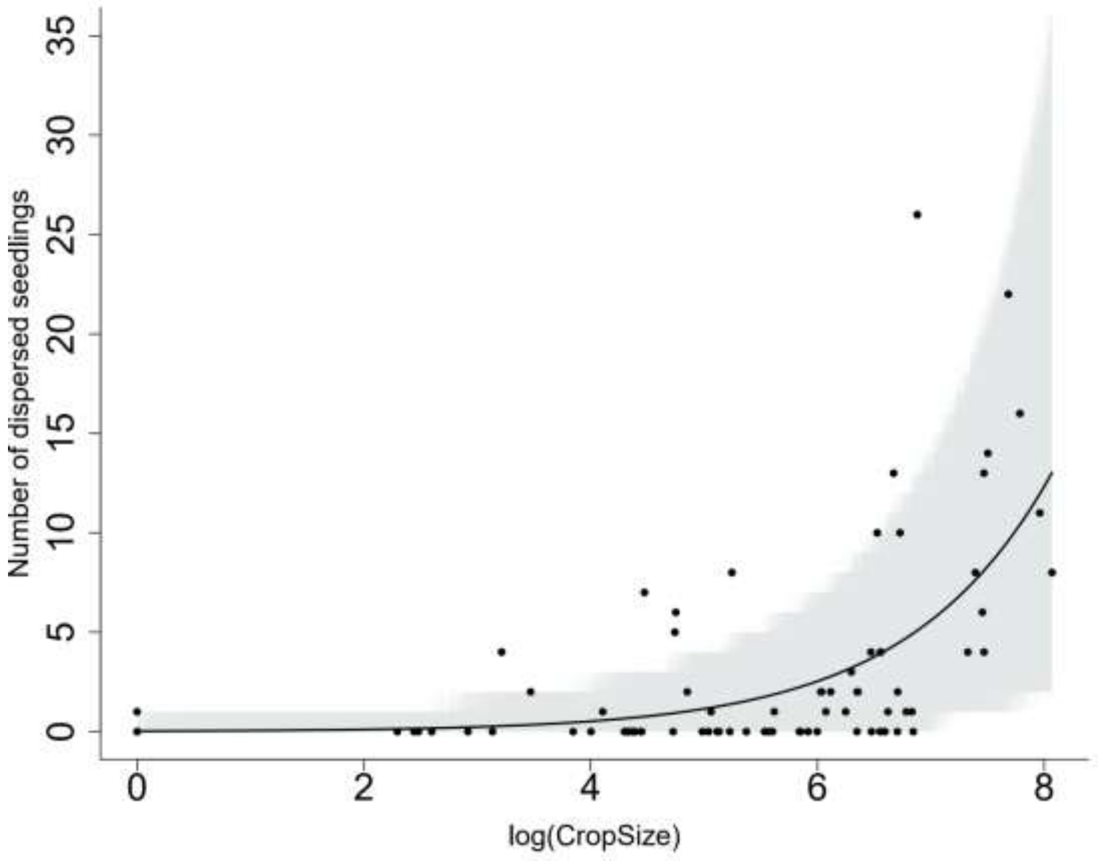
